# Chronic Alcohol Exposure Alters Expression of GluN3A-NMDA Receptors in Adult VTA DA Neurons

**DOI:** 10.64898/2026.09.08.750197

**Authors:** Kechun Yang, Jessica K. Shaw, John A. Dani, Mariella De Biasi

## Abstract

N-methyl-D-aspartate receptor (NMDAR) neurotransmission plays a central role in the neurophysiological and behavioral adaptations produced by chronic alcohol (EtOH) exposure and withdrawal. Although alcohol-induced changes in canonical NMDAR subtypes have been extensively studied, the mechanisms by which prolonged alcohol exposure remodels NMDAR function remain incompletely understood. To investigate the impact of chronic voluntary alcohol consumption on glutamatergic signaling, adult C57BL/6J mice were exposed to 20% EtOH using the intermittent two-bottle choice (I2BC) approach for 8 weeks and NMDAR function was examined in ventral tegmental area (VTA) dopamine (DA) neurons using whole-cell electrophysiology. Chronic EtOH exposure reduced NMDA-evoked currents, an effect that became more pronounced during withdrawal. Withdrawal also reduced NMDAR outward current rectification, indicating that chronic alcohol exposure altered the biophysical properties of NMDAR-mediated signaling. These unexpected observations led us to investigate the contribution of GluN3A-containing NMDARs. Pharmacological analyses revealed a significant increase in the proportion of VTA DA neurons expressing functional GluN3A-containing receptors following chronic EtOH exposure, with all tested neurons expressing these receptors during withdrawal. Glycine activation directly depolarized VTA DA neurons and increased action potential firing, demonstrating that the recruited receptors function as excitatory NMDARs. Moreover, the alcohol-induced changes in NMDAR currents, receptor pharmacology, and glycine responsiveness were absent in GluN3A null mice. Together, these findings identify functional recruitment of GluN3A-containing NMDARs as a previously unrecognized neuroadaptation through which chronic alcohol remodels glutamatergic signaling in the adult VTA.

## INTRODUCTION

Alcohol use disorder (AUD) remains a major public health challenge and is among the leading causes of preventable morbidity and mortality worldwide, accounting for 4.7% of all global deaths^1–3^. Chronic alcohol exposure induces widespread neuroadaptations throughout the central nervous system that contribute to the behavioral manifestations of AUD^4–9^. A large body of evidence from studies in humans^10,11^, non-human primates^12^, and rodents^13–20^ demonstrates that ethanol alters glutamatergic transmission including within the mesocorticolimbic dopamine (DA) system that originates in the ventral tegmental area (VTA) and comprises the canonical “reward system”. Chronic alcohol exposure also disrupts glutamate homeostasis and alters the expression, localization, and function of glutamate receptors throughout reward-related brain circuits^20–24^ .

DA neurons within the VTA integrate excitatory and inhibitory inputs to regulate reinforcement, reward prediction, motivation, and behavioral responses to drugs of abuse^25–30^. Both glutamatergic afferents to the VTA and glutamatergic neurons residing within the VTA contribute to the regulation of DA neuron activity and undergo significant neuroadaptations following chronic alcohol exposure and withdrawal^15,16,25–27^. Among ionotropic glutamate receptors, N-methyl-D-aspartate receptors (NMDARs) are particularly important because they are both acute targets of ethanol and key mediators of long-lasting, alcohol-induced neuroplasticity. Consequently, they have been implicated in intoxication, dependence, withdrawal, craving, and relapse^4,16,31–36^.

NMDARs are tetrameric receptors assembled from two obligatory GluN1 subunits together with GluN2 (GluN2A-D) and/or GluN3 (GluN3A-B) subunits^37–39^. Their subunit composition determines receptor kinetics, Ca²⁺ permeability, voltage dependence, and downstream intracellular signaling, thereby exerting a major influence on synaptic plasticity. Consequently, considerable effort has focused on understanding how alcohol regulates NMDARs, especially those containing GluN2A and GluN2B subunits. Chronic alcohol exposure and withdrawal alter the expression and function of these receptor subtypes in several brain regions, where they contribute to alcohol-related neuroadaptations and drinking behaviors^40–48^. In contrast, the potential contribution of GluN3-containing NMDARs to alcohol-induced plasticity remains largely unexplored.

GluN3A is a particularly intriguing NMDAR subunit because its physiological role differs fundamentally from that of the canonical GluN2 subunits. Incorporation of GluN3A reduces Ca²⁺ permeability, decreases voltage-dependent Mg²⁺ block, and modifies receptor gating, thereby fundamentally altering the biophysical properties and signaling characteristics of NMDAR-mediated transmission^37–39,49,50^.

During early postnatal development, GluN3A is widely expressed throughout the brain, where it contributes to synaptic maturation by limiting NMDAR-dependent plasticity^51–55^. Although GluN3A expression significantly declines as neural circuits mature^49–51^, functional GluN3A-containing receptors persist in selected adult brain regions—including the VTA—where their physiological roles remain incompletely understood^56–61^. These receptors may exist either as glycine-gated GluN1/GluN3A di-heteromers or as glutamate/glycine-gated tri-heteromeric receptors containing GluN1, GluN2, and _GluN3A56,57,59-62._

The developmental regulation and unique physiological properties of GluN3A suggest that functional recruitment of GluN3A-NMDARs in the adult brain could provide a unique mechanism for remodeling NMDAR signaling and synaptic function. Indeed, transient recruitment of GluN3A-containing NMDARs is required for cocaine-induced synaptic plasticity in VTA DA neurons^61^, raising the possibility that GluN3A recruitment may represent a common mechanism through which addictive drugs remodel mature reward circuits. Whether chronic alcohol exposure similarly induces functional expression of GluN3A-containing NMDARs in the adult VTA is not presently known.

To address this question, we exposed adult mice to chronic voluntary alcohol consumption using the intermittent two-bottle choice (I2BC) paradigm^63,64^ and examined NMDAR function in identified VTA DA neurons using whole-cell electrophysiology. Our findings demonstrate that chronic alcohol exposure and withdrawal recruit functional GluN3A-containing NMDARs in adult VTA dopamine neurons, identifying a previously unrecognized mechanism through which alcohol remodels glutamatergic signaling in the mesolimbic reward system.

## EXPERIMENTAL PROCEDURES

### Animals

Adult C57Bl/6J (The Jackson laboratory, strain # 000664) and GluN3A null (GluN3A -/-; The Jackson laboratory, B6;129X1-*Grin3atm1Nnk*/J strain # 029974) mice of both sexes were bred and raised in house. All mice were single-housed at the start of the experiments in standard “shoebox” cages (7.6 in × 15 in × 5.1 in) in a temperature- and humidity-controlled facility under a reverse 12-h light/dark cycle. They had ad libitum access to food and fluid. All procedures were carried out during the dark cycle and in compliance with the approved guidelines specified by the Institutional Animal Care and Use Committee at the University of Pennsylvania.

### Chronic Ethanol Treatment

To model chronic, voluntary exposure to ethanol (EtOH), mice were exposed to EtOH using the I2BC drinking paradigm (Fig. 1A) as described previously^63,64^. Mice were first habituated to the two-bottle setup for 1 week prior to the initiation of chronic EtOH exposure. EtOH was administered using 50-ml Nunc^TM^ conical centrifuge tubes (Thermo Fisher Scientific, Waltham, MA) with one-hole rubber stoppers with straight stainless-steel, open-tip sippers. The I2BC consisted of intermittent access to EtOH over three, 24-h sessions/week on Mondays, Wednesdays, and Fridays. Mice were first exposed to a one-week period of acclimation, during which the concentration of EtOH increased with each presentation (3, 6, and 10% v/v), followed by a maintenance phase with ≥8 weeks of intermittent access to 20% EtOH (Fig. 1A). On the remaining days of the week, mice had access to two bottles of water. Bottle positions were switched for each EtOH session to avoid side preference. EtOH-treated mice were then randomly assigned to be either EtOH-sated (Sated) or EtOH-withdrawn (WD). Twenty-four hours before the day of patch clamp recording, mice in the Sated group received EtOH as the sole source of fluid while WD mice received only water (Fig 1A). This approach was adopted to decrease variability among subjects within each group. Control mice had access only to water throughout the duration of the experiments.

**Figure 1.**
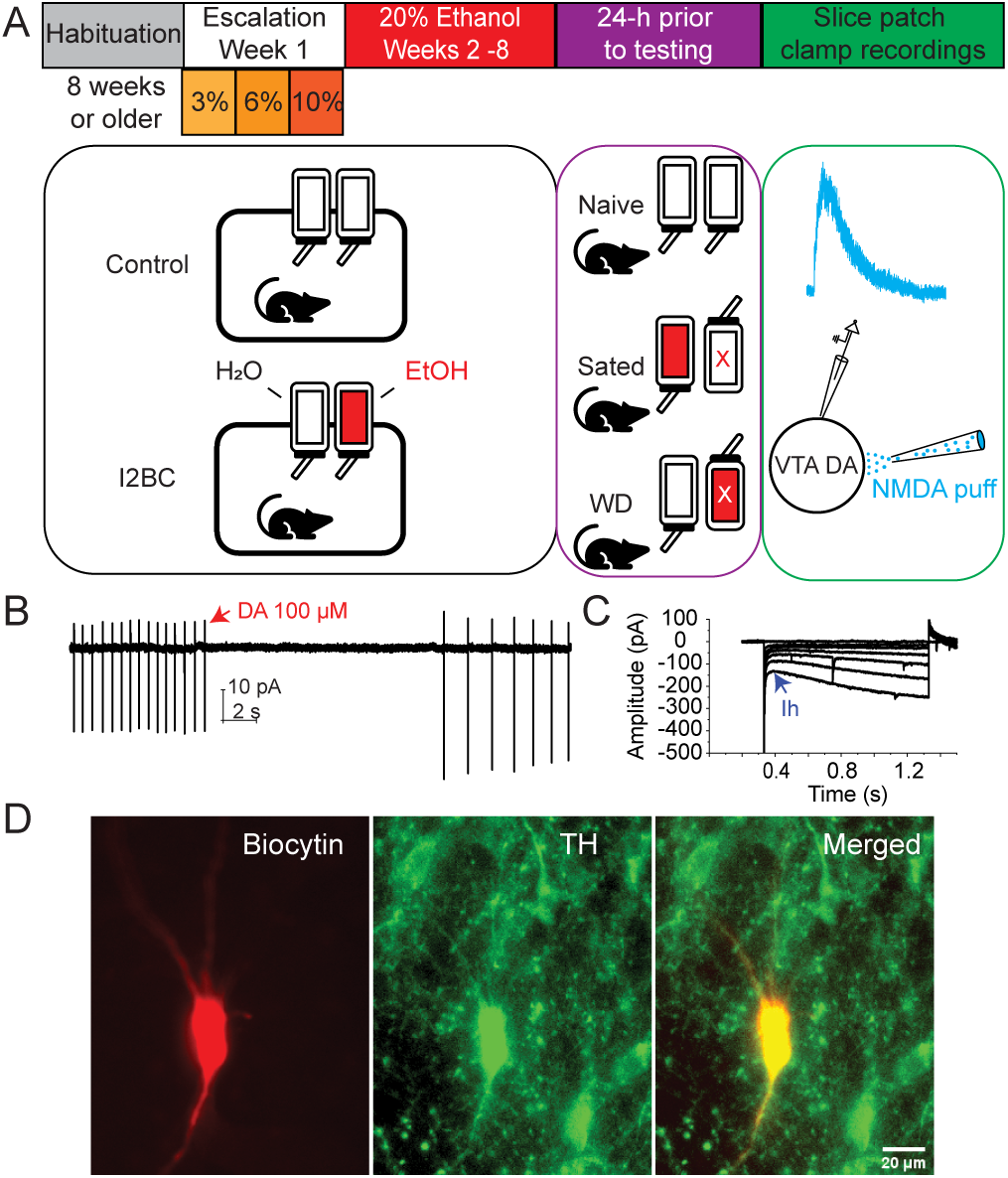
Schematic of chronic Intermittent Two-bottle choice (I2BC) EtOH drinking and patch clamp recordings in VTA DA neurons. **A.** An illustration of the timeline for chronic I2BC EtOH drinking with mice 8-weeks or older. Following habituation to the facility/single-housing, mice receive increasing concentrations of EtOH over 1 week under an I2BC schedule followed by ≥8w maintenance at 20%. 24 hr prior to the electrophysiology recordings (right panel), Sated mice receive only EtOH and withdrawal (WD) mice receive only water; Naïve mice receive only water throughout. **B.** Indicative of a DA neuron, spontaneous action potential firing was transiently stopped by a puff-application of 100 µM DA. **C**. Indicative of a DA neuron, hyperpolarization-activated cationic currents (Ih) were evoked by hyperpolarizing pulses. **D**. Example of a biocytin-filled VTA cell confirmed to be a dopaminergic neuron using TH immunohistochemical staining.

### Slice Preparation

Mice were euthanized with a 0.2 ml intraperitoneal injection containing 16 mg ketamine + 4 mg xylazine followed by exsanguination via transcardial perfusion, performed as previously described^65–67^ using ice-cold N-methyl-D-glucamine (NMDG)-based artificial cerebrospinal fluid (ACSF, in mM): 92 NMDG, 2.5 KCl, 1.2 NaH_2_PO_4_, 30 NaHCO_3_, 20 HEPES, 25 glucose, 2 thiourea, 5 Na-ascorbate, 3 Na-pyruvate, 0.5 CaCl_2_, and 10 MgSO_4_, pH 7.3-7.4 with concentrated HCl. After decapitation, the brain was rapidly extracted and sectioned using a Leica VT1200S vibratome in ice-cold, well-oxygenated NMDG ACSF. Horizontal slices containing the VTA (230 µm) were immediately transferred to oxygenated NMDG ACSF incubated in a thermostatic water bath (Precision Water Bath Model 180, Winchester, VA) set at 32°C for about 13 min. The slices were further recovered in HEPES-based holding ACSF (in mM): 92 NaCl, 2.5 KCl, 1.2 NaH_2_PO_4_, 30 NaHCO_3_, 20 HEPES, 25 glucose, 2 thiourea, 5 Na-ascorbate, 3 Na-pyruvate, 2 CaCl_2_, and 2 MgSO_4_ at room temperature for at least 1-h prior to recording.

### Patch-clamp Recordings

Slice recordings were performed in a modified Quick Exchange Recording Chamber (Warner Instruments, Hamden, CT) that continuously perfused the slice with well-oxygenated standard recording ACSF (in mM): 124 NaCl, 2.5 KCl, 1.2 NaH_2_PO_4_, 24 NaHCO_3_, 5 HEPES, 12.5 glucose, 2 CaCl_2_, and 2 MgSO_4_. The ACSF recording solution was maintained at 32-34℃ using an inline heater system (TC-324B, Warner Instruments, Hamden, CT). Thin wall WPI borosilicate glass capillaries (TW150-4, 1.12 mm ID, 1.5 mm OD; Sarasota, FL) were used to pull recording and puff electrodes with resistances of 2-3 MΩ after filling with internal solution, utilizing a Narishige Dual Stage Puller (PC-10, Tokyo, Japan). Putative DA neurons in the VTA were visually identified with the aid of a Leica DM 6000 FS fluorescence microscope and patch-clamp recordings were then conducted using MutiClamp 700B and Axon Digidata 1550 (Molecular Devices). Responses to NMDA/glycine puffs were recorded using a cesium methanesulfonate-based internal solution (in mM): 130 Cs-methanesulfonate, 5 QX-314, 10 HEPES, 5 Phosphocreatine, 0.1 Na_4_-EGTA, 10 TEA-Cl, 3.8 NaCl, 0.4 Na GTP, 2 Mg ATP, 1 MgSO4 (pH 7.2, 280–290 mOsm), used to measure NMDA-currents at +60 mV and to further examine I-V relationships where the holding voltage varied from +60 mV to -60 mV. The rest of the studies employed a K-gluconate-based intracellular solution (in mM): 140 K-gluconate, 5 KCl, 10 HEPES, 0.2 EGTA, 2 MgCl_2_, 4 MgATP, 0.3 Na_2_GTP, and 10 Na_2_-phosphocreatine (pH 7.3 with KOH). All recordings started with the tight-seal patch-clamp recording configuration^65,67^ to obtain spontaneous action potential firing in most recorded neurons under voltage clamp mode with the holding potential set at 0 mV^68^. DA neurons were first identified based on their firing patterns and the sensitivity to puff-applied DA (100 µM) (Fig. 1B). The whole-cell patch-clamp configuration was then achieved by application of transient strong suctions. Recorded neurons were further confirmed as DA neurons by the presence of the hyperpolarization-activated cationic currents (H-current^65–67^, Fig. 1C). A subset of the recorded neurons was labeled intracellularly with biocytin through the recording electrode to confirm them as DA neurons using tyrosine hydroxylase (TH) staining (Fig. 1D). Either NMDA or glycine was pressure-puffed locally through a patch pipette (Fig. 1A panel, lower right) by a Picospritzer II (Parker Instrumentation, Fairfield, NJ) every 2 min in the ACSF bath containing either 100 µM picrotoxin only or 100 µM picrotoxin and 50 µM strychnine (to inhibit GABA and glycine receptors, respectively)^56,59,60^. The puffing pipette was typically positioned around 20 to 40 µm away from the soma of the recorded DA neurons. Only recordings with access resistance (Ra) < 10 MΩ were accepted for further analysis using the analysis program, Clampfit.

### Immunohistochemistry

After filling recorded neurons with biocytin during recording, the slices were immediately fixed overnight in 4% paraformaldehyde at + 4°C and were washed with PBS, followed by heat-mediated (30 minutes at 45°C) antigen retrieval (10 mM sodium citrate, pH 8.5). Slices were then incubated in 50 mM glycine at room temperature for about 2 hours. Following another rinse with PBS, slices were incubated in a blocking solution comprising 5% bovine serum albumin (BSA, Sigma), 5% normal goat serum (NGS, Vector Laboratories), and 0.5% Triton-X (VWR) at room temperature for 3 hours. Afterwards, slices were incubated overnight at 4°C in a primary antibody solution containing 1xPBS, 1% BSA, 1% NGS, 0.1% Triton-X, and rabbit anti-tyrosine hydroxylase (1:1000, Abcam, #ab112). The following day, slices were first rinsed and then incubated for 7 hours at room temperature in a secondary antibody solution containing PBS, 1% BSA, 1% NGS, 0.1% Triton X, streptavidin Alexa Fluor 568 (1:2000; Thermo Fisher Scientific, #S11226), and goat anti-rabbit Alexa Fluor 488 (1:100, Thermo Fisher Scientific, #A-11008). After 3 final rinses in 0.1M sodium phosphate buffer (pH 7.35), slices were mounted on glass slides (Superfrost Plus, Fisher), coverslipped with fluorescent mounting medium (Fluoromount-G, Thermo Fisher Scientific), and were imaged on an epifluorescence microscope to confirm TH expression in recorded neurons (Fig. 1D).

### Data Analysis

All values are reported as Mean ± SEM. Statistical analyses were conducted using SPSS (v27), OriginPro 2024, and R. Comparisons between two groups were performed using a Student’s t-test (OriginPro). Analyses comparing more than two groups were conducted using one-way or two-way repeated measures ANOVAs as appropriate (SPSS), with Sidak’s correction for multiple comparisons and Greenhouse-Geisser corrections for sphericity. The non-parametric Kruskal-Wallis test was used for non-normally distributed data, with follow-up pairwise comparisons with Bonferroni adjusted p-values. Separate Firth’s logistic regressions were used to compare proportions of cells responsive to glycine with and without bath application of CGP using the logistf package (R).

### Drugs

5,7-dichlorokynurenic acid, sodium salt (5,7-DCKA) was purchased from EMD (Millipore Corp, USA). Dopamine (DA), strychnine, and (3-chlorophenyl)(6,7-dimethoxy-1-((4-methoxyphenoxy)methyl)-3,4-dihydroisoquinolin-2(1 H)-yl)methanone (CIQ) were purchased from Sigma-Aldrich. All other chemicals were procured from Tocris Bioscience. NMDA, glycine, and DA were locally puffed onto the recorded neurons while CGP-78608 ((1S)-1-[[(7-Bromo-1,2,3,4-tetrahydro-2,3-dioxo-5-quinoxal-inyl)methyl]amino]ethyl] phosphonic acid), DL-APV (DL-2-Amino-5-phosphonopentanoic acid), and 5,7-DCKA were continuously bath applied.

## RESULTS

### Chronic alcohol exposure and withdrawal suppress canonical NMDAR signaling in VTA dopamine neurons

Altered glutamatergic function is key to the pathophysiology of EtOH seeking^5,9^, and VTA DA neurons are among the targets of the neuroadaptations associated with chronic alcohol consumption^5^. Surprisingly, there is little information available on the effects of chronic EtOH on NMDAR function in the VTA. To investigate potential changes in NMDAR function after chronic EtOH exposure, we measured NMDAR-mediated whole-cell currents evoked by puff-applied 100 µM NMDA onto VTA DA neurons from mice voluntarily drinking 20% EtOH in the I2BC paradigm (Fig. 2A). Chronic EtOH significantly reduced the amplitude of NMDAR currents [Fig. 2B, Kruskal-Wallis H(2)=31.648, p<0.001]. Peak current was reduced from 1351.6 ± 90.4 pA in EtOH naïve controls (Fig. 2B, naïve) to 712.6 ± 62.5 pA in EtOH sated mice (Fig. 2B, sated; p<0.01). A further reduction in NMDAR currents was recorded in DA neurons from mice 24 hours after withdrawal from chronic EtOH (Fig. 2B, WD, 374.5 ± 89.9, p<0.01), but the difference between EtOH-sated and WD was not statistically significant.

**Figure 2.**
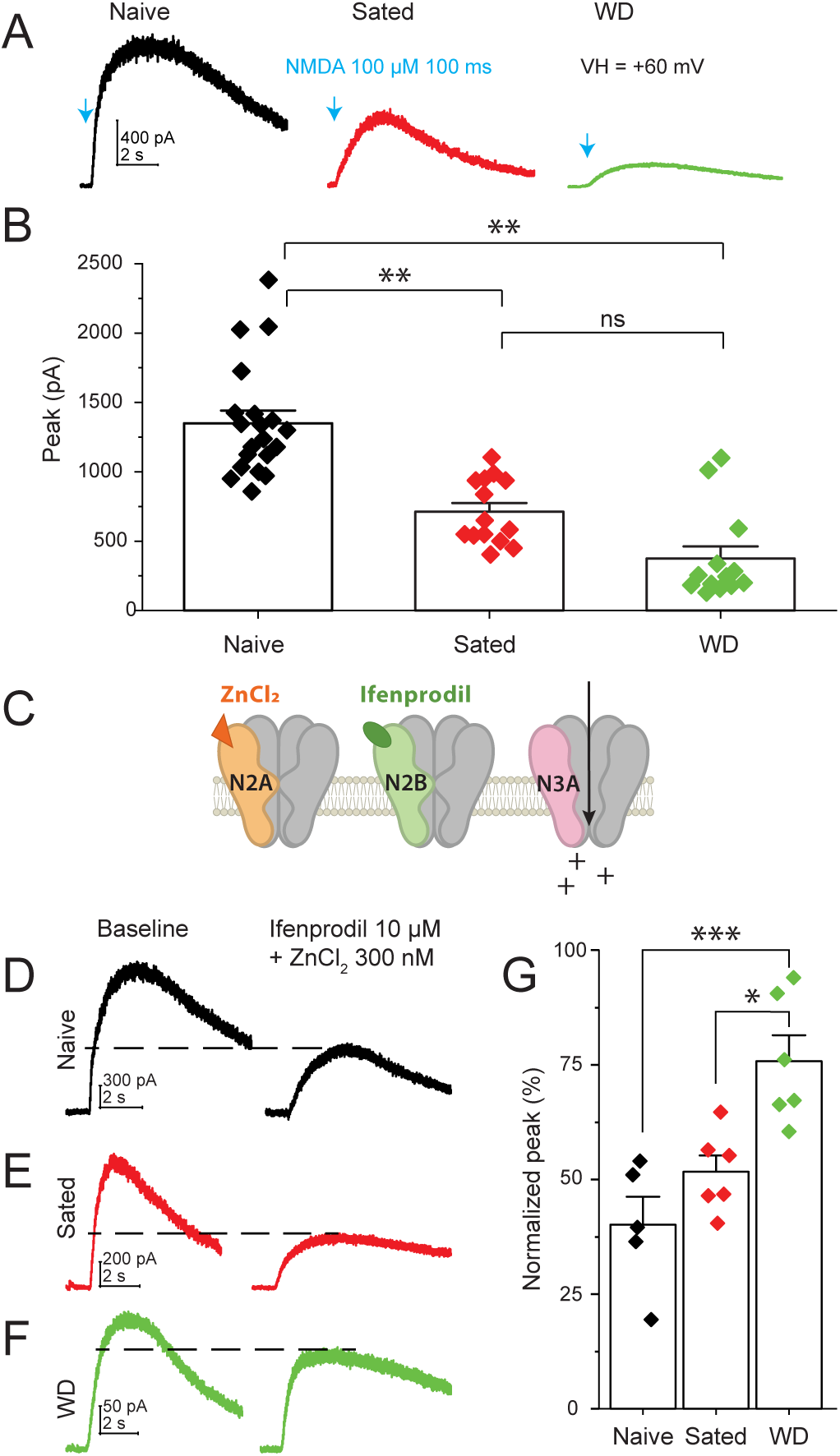
Suppressed responses to NMDA and reduced GluN2 inhibition after chronic EtOH and WD in VTA DA neurons. **A.** Representative traces of whole-cell currents induced by puff-application of 100 µM NMDA onto VTA DA neurons at baseline (EtOH naïve, left), during chronic EtOH (Sated, middle), and during EtOH withdrawal (WD, right). Blue arrows indicate the puff application of NMDA. The recorded neurons were clamped at +60 mV. **B**. Summary of the peak amplitude of NMDA-induced currents. Chronic EtOH exposure and WD significantly suppressed NMDA currents. Neurons from both the sated (n=14 cells from 4 males & 4 females) and WD (n=13 cells from 5 males & 4 females) groups significantly differed from those from EtOH-naïve mice (n=20 cells from 5 males & 7 females). However, there was no significant difference between neurons from the sated and WD animals (p=0.299). **C.** Cartoon depicting the use of ZnCl_2_ and Ifenprodil to suppress GluN2-mediated responses to NMDA. ZnCl_2_ preferentially inhibits GluN2A-containing while Ifenprodil inhibits GluN2B-containing NMDARs. **D-F**. Typical NMDA currents before (left, baseline) and after (right) bath application of 10 µM Ifenprodil and 300 nM ZnCl_2_ in VTA DA neurons in naïve (D), sated (E), and WD (F) mice. **G.** Summary of the normalized inhibitory effects of Ifenprodil and ZnCl_2_ on NMDA currents. All data collected after bath application of the antagonists were normalized to their baseline (100%). NMDA currents were inhibited by Ifenprodil and ZnCl_2_ to 40.0 ± 6.1% (Naïve, n=5 cells from 2 males & 2 females), 51.7 ± 3.6% (Sated, n=6 cells from 3 males & 2 females), and 75.8 ± 5.6% (WD, p<0.01, n=6 cells from 2 males & 3 females), respectively. One-way ANOVA detected significant differences in suppression by Ifenprodil and ZnCl_2_ across the 3 groups of mice p<0.001) and Sidak’s post-hoc further confirmed that NMDA currents from EtOH WD mice were less sensitive to Ifenprodil and ZnCl_2_ compared to both naïve and EtOH sated mice (*** p<0.001, ** p≤0.01, * p<0.05, ns: not significant).

### The functional contribution of GluN2A/GluN2B-containing NMDARs is reduced during alcohol exposure and withdrawal

The dramatic changes in the amplitude of NMDAR currents observed in alcohol-exposed mice might reflect changes in NMDAR subunit composition. As chronic alcohol has been reported to alter the expression and function of GluN2A- and GluN2B-containing NMDARs in multiple brain regions outside of the VTA^40–45^, we next asked whether the reduction in NMDA-evoked currents we observed in VTA DA neurons was due to changes in these canonical receptor subtypes. We therefore measured the sensitivity of NMDA currents to the GluN2B antagonist ifenprodil and the GluN2A antagonist ZnCl₂ across alcohol-naïve, alcohol-sated, and withdrawn mice (Fig. 2C-F). Bath application of a combination of Ifenprodil (10 µM), and ZnCl_2_ (300 nM)^61^, led to significantly different inhibition of NMDAR currents recorded in slices from naïve vs. sated and WD mice (Fig. 2G; one-way ANOVA [F(2,14)=12.481, p<0.001]). In EtOH-naïve mice, the combined antagonists inhibited approximately 60% of the NMDA-gated current (Fig. 2D,G; residual current = 40.0 ± 6.1 % of baseline), consistent with GluN2A- and GluN2B-containing receptors accounting for the majority of NMDAR-mediated signaling under basal conditions in the brain^69,70^. In EtOH-sated mice, the antagonist-sensitive component was smaller (Fig. 2E,G; residual current = 51.7 ± 3.6 % of baseline), although this decrease did not reach statistical significance. In WD mice, however, inhibition by ifenprodil and ZnCl_2_ was markedly reduced, with only ∼30% of the NMDA current blocked (Fig. 2F,G; residual current = 71.6± 5.6% of baseline; p<0.001 vs naïve, p<0.05 vs sated). Overall, these findings indicate that GluN2A- and GluN2B-containing receptors account for the majority of functional NMDAR signaling in alcohol-naïve VTA dopamine neurons (Figure 2D, black), but that their contribution declines following chronic alcohol exposure (Figure 2E, red) and is further diminished during withdrawal (Figure 2F, green). The reduced sensitivity to GluN2A/GluN2B antagonism suggests that a growing fraction of the remaining NMDA current is mediated by receptors with a different subunit composition.

To further investigate the identity of the receptors contributing to the NMDA current after chronic EtOH exposure, we next examined the current-voltage (I-V) relationship of NMDA-evoked currents in VTA DA neurons from EtOH-naïve, EtOH-sated, and EtOH-withdrawn mice (Fig. 3A). A two-way mixed-design ANOVA revealed a significant interaction between membrane potential and ethanol exposure [Fig.3B, F(3.83, 26.811) =3.693, p<0.05]. Pairwise comparisons showed that NMDA-gated currents recorded from WD mice exhibited significantly reduced outward rectification (Fig. 3B, green triangles) compared to both EtOH naïve (Fig. 3B, black squares) and EtOH sated (Fig. 3B, red circles) mice (p≤0.01 between -60 mV and 0 mV). This altered voltage dependence is consistent with reduced Mg²⁺ block at hyperpolarized membrane potentials of inward going cationic current. This change in Mg^2+^ block during withdrawal is consistent with a change in NMDARs’ subunit composition. Together with the reduction in GluN2A/GluN2B-mediated currents, these findings suggested that chronic EtOH exposure recruits a distinct population of functional NMDARs with biophysical properties consistent with GluN3A-containing receptors NMDARs^49,51,61^. We tested this possibility directly in the following experiments.

**Figure 3.**
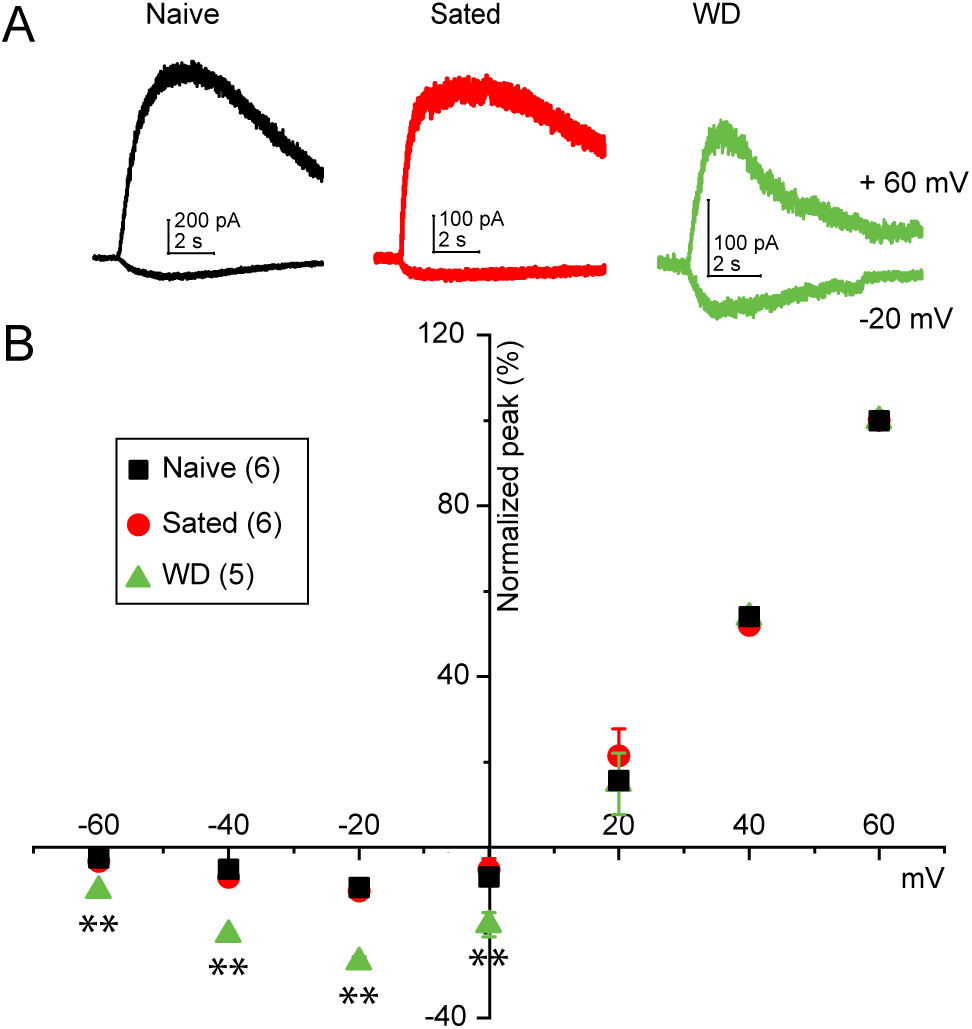
Withdrawal from chronic EtOH exposure alters the I-V relationships of NMDA currents in WT mice. **A**. Typical responses to puff-applied 100 µM NMDA while recorded VTA DA neurons were clamped at either +60 mV (top) or -20 mV (bottom) in naïve (left), EtOH-sated (middle), and WD (right) WT mice. **B.** Averaged, normalized I-V curves of NMDA currents across different EtOH states. A two-way mixed-design ANOVA revealed significant interactions (F(3.830, 26.811) = 3.693, p<0.05) between EtOH state and the clamped voltage (mV). Pairwise comparisons with Sidak’s adjustment for multiple comparisons showed that WD (n=5 cells from 3 males & 2 females) was significantly different from naïve (n=6 cells from 2 males & 3 females) and EtOH sated (n=6 cells from 2 males & 2 females) mice between -60mV and 0mV. All data were normalized to the values obtained at +60 mV (100%). ** p≤0.01.

### Chronic EtOH exposure and withdrawal promote functional recruitment of GluN3A-NMDARs in VTA dopamine neurons

The pharmacological and biophysical properties of the NMDAR currents described above suggested that chronic EtOH exposure recruits GluN3A-NMDARs in adult VTA DA neurons. To test this hypothesis directly, we took advantage of the unique property of GluN3A-NMDARs whereby glycine can activate the receptor through binding to the GluN3A subunit. Therefore, to maximize the detection of GluN3A-mediated responses, we locally applied glycine puffs (1 mM) while inhibitory glycine receptors were blocked with strychnine. An important consideration is that glycine also binds to the GluN1 subunit, where it acts as a co-agonist and accelerates receptor desensitization^71,72^. Therefore, to maximize detection of GluN3A-mediated currents, recordings were repeated following bath application of 500 nM CGP-78608, a highly selective GluN1 competitive antagonist that relieves glycine-induced receptor desensitization and unmasks GluN3A-mediated responses^59,73,74^. This strategy allowed us to compare the prevalence of functional glycine-responsive GluN3A-containing receptors across the different EtOH states (Fig. 4). In EtOH naïve mice, only 7.7% (2/26) of the recorded VTA DA neurons responded to glycine in the absence of CGP-78608 (Fig. 4 A). Following incubation with CGP-78608, the proportion of glycine-responsive neurons increased to 43.8% (7/16), indicating that functional GluN3A-containing receptors are present in a subset of adult VTA DA neurons even under basal, EtOH-naïve conditions (Fig 4A). In EtOH sated mice, 25% (5/20) of recorded DA neurons responded to glycine in the absence of CGP-78608 and the proportion of glycine-responsive neurons increased to 58.8% (10/17) after receptor desensitization was blocked with CGP-78608 (Fig. 4B), but these proportions were not significantly different from EtOH-naïve mice. The most striking differences were observed during withdrawal. Neurons from EtOH-withdrawn mice were significantly more likely to display glycine-gated currents than EtOH-naïve cells (OR=33.8, 95% CI: 7.3-225, p<0.0001; Fig 4C,D), with 78.9% (15/19) of recorded neurons displaying glycine-gated currents even in the absence of CGP-78608. After CGP-78608 application, all recorded neurons (16/16) were responsive to glycine (OR=48.1, 95% CI: 4.3-5657.5, p<0.001; Fig. 4C,E). Together, these observations demonstrate that functional GluN3A-containing NMDARs are present in a small subset of adult VTA DA neurons under basal, EtOH-naive conditions and are recruited following chronic EtOH exposure, becoming nearly ubiquitous during withdrawal.

**Figure 4.**
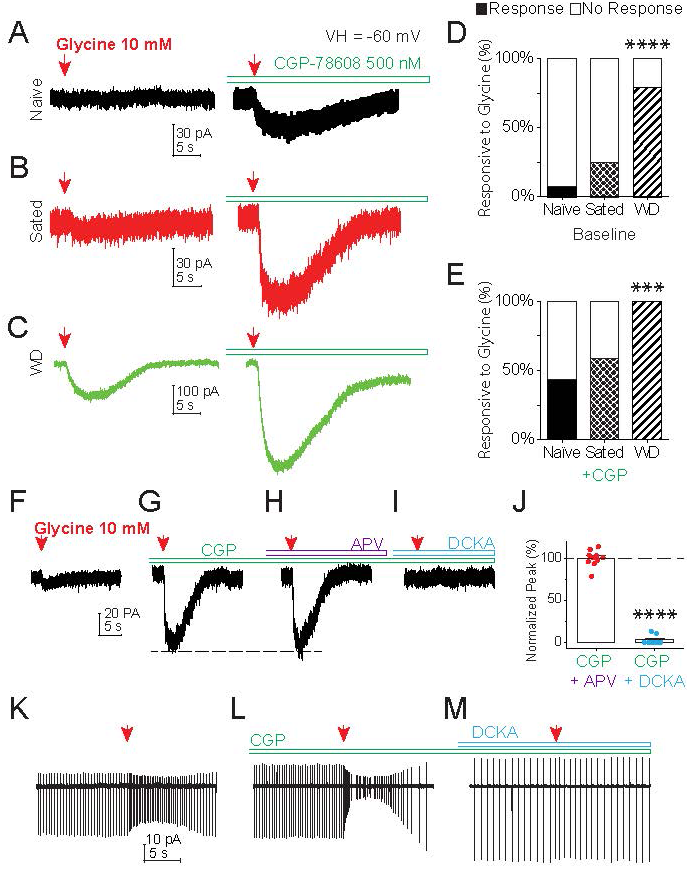
Withdrawal from chronic EtOH exposure increases the probability of glycine-gated GluN3A-mediated currents recorded in WT VTA DA neurons. **A-C**. Whole-cell currents induced by puff application of 10 mM glycine in VTA DA neurons from either naïve (A), sated (B), or WD (C) mice before (left) and after (right) bath application of a GluN1 inhibitor (CGP-78608, 500 nM, indicated by the left side opened bars), in the presence of 50 µM strychnine to block the inhibitory glycine receptors. **D-E**. Summaries of the proportions of glycine currents recorded in VTA DA neurons before (baseline, D) and after (+ CGP, E) the application of CGP in naïve (26 cells from 10 males & 8 females), sated (20 cells from 7 males & 8 females), and WD (19 cells from 7 males & 7 females) mice. The probability of a response to glycine was significantly greater in WD mice than naïve under both conditions. **F-I**. Currents in response to 10 mM glycine puff-applied every 2 min (F) during bath application of 500 nM CGP alone (G), CGP and 100 µM APV (H), and CGP with 20 µM 5, 7-DCKA (I) recorded in a VTA DA neuron from WT mice undergoing withdrawal from chronic EtOH exposure. **J.** Summary of the effects of APV and DCKA on the currents induced by glycine. All data (from 4 males & 4 females) were normalized to the baseline indicated by a dashed line (100%, corresponding to the currents recorded after bath application of CGP). Glycine-induced currents were significantly lower after DCKA (paired t-test, p<0.0001) but not APV. **K-M.** Spontaneous action potential firing in response to puff-applied glycine alone (K), after continuous bath application of CGP (L), and CGP plus DCKA (M). Left side opened bars indicate continuous bath application of the antagonists (n= 6 cells from 3 males & 1 female). ***p<0.001, ****p<0.0001

Demonstrating glycine-sensitive currents establishes the presence of functional GluN3A-containing receptors but does not determine whether these receptors contribute to neuronal excitability. We therefore next asked whether activation of GluN3A-containing NMDARs could directly influence VTA DA neuron activity. We first investigated the pharmacological properties of the glycine-induced currents. Glycine alone elicited small excitatory currents in recorded VTA DA neurons from EtOH-withdrawn mice even in the absence of CGP-78608 (Fig. 4F), and these responses were greatly potentiated following bath application of 500 nM CGP-78608 to slow receptor desensitization (Fig. 4G), consistent with relief of GluN1-mediated receptor desensitization. The glycine-induced currents were unaffected by 100 µM APV, a glutamate binding-site NMDAR antagonist (Fig. 4H,J; 99.94 ± 3.09% of currents after CGP-78608 incubation alone) but were almost completely blocked by 20 µM 5,7-DCKA, a GluN1/GluN3A glycine-binding site antagonist (Fig. 4I,J; 2.59 ± 1.72% of the currents measured after CGP-78608, paired t-test, p<0.0001). Together, the potentiation of glycine-evoked currents by CGP-78608, their near-complete blockade by DCKA, and their insensitivity to APV are the pharmacological hallmarks of GluN3A-containing NMDARs^59,60^ and confirm that the glycine-induced currents were mediated by GluN3A-containing receptors.

We next determined whether these receptors were operational and capable of modulating neuronal activity. Using the tight-seal, cell-attached, patch-clamp mode, spontaneous action potential firings were recorded in VTA DA neurons from mice 24 h after EtOH withdrawal. Glycine puff application significantly increased spontaneous firing rate from 1.9 ± 0.77 Hz to 3.5 ± 1.93 Hz (Fig. 4K, paired t-test, p<0.05). Bath application of 500 nM CGP-78608 further enhanced the excitatory effect of glycine, producing sustained depolarization that culminated in depolarization block (Fig. 4L). Conversely, blocking the glycine-binding site with 20 µM 5,7-DCKA abolished the excitatory effect of glycine and restored spontaneous firing to a steady level (Fig. 4M). Together, these findings demonstrate that the GluN3A-containing NMDARs recruited following chronic EtOH exposure are fully operational excitatory receptors capable of driving membrane depolarization and increasing action potential firing in adult VTA dopamine neurons during withdrawal.

### GluN3A is required for alcohol-induced remodeling of NMDAR function in VTA dopamine neurons

To determine whether GluN3A is required for the alcohol-induced NMDAR neuroadaptations observed in VTA DA neurons, we repeated our electrophysiological analyses in GluN3A knockout (GluN3A-/-) mice undergoing the same I2BC drinking paradigm. Unlike what we observed in control mice, chronic EtOH exposure and withdrawal failed to reduce NMDA-evoked currents in GluN3A-deficient mice (Fig. 5A,B). NMDA current amplitudes in response to 100 µM NMDA puffs were similar in EtOH-naïve (493.2 ± 48.5 pA), EtOH-sated (398.7 ± 133.7 pA), and EtOH-withdrawn mice (406.9 ± 82.9 pA) when analyzed using one-way ANOVA [F(2,35)=0.91, p=0.41], in strong contrast with the reduction in NMDA currents observed in WT mice (Fig. 2). Consistent with the absence of GluN3A, glycine failed to evoke detectable currents in any recorded VTA DA neuron, even after bath application of 500 nM CGP-78608 (Fig. 5C). Likewise, chronic EtOH exposure failed to alter the voltage dependence of NMDA-evoked currents in GluN3A-/-mice. Current-voltage relationships were indistinguishable among EtOH-naïve, EtOH-sated, and EtOH-withdrawn animals (Fig. 5D), with no significant effect of EtOH state [two-way mixed-design ANOVA, F(2,34)=2.176, p=0.129] and no interaction between EtOH state and membrane potential [F(3.194,54.301)=1.704, p=0.174]. Thus, the reduction in outward rectification observed following EtOH withdrawal in WT mice was completely absent in GluN3A-deficient animals.

**Figure 5.**
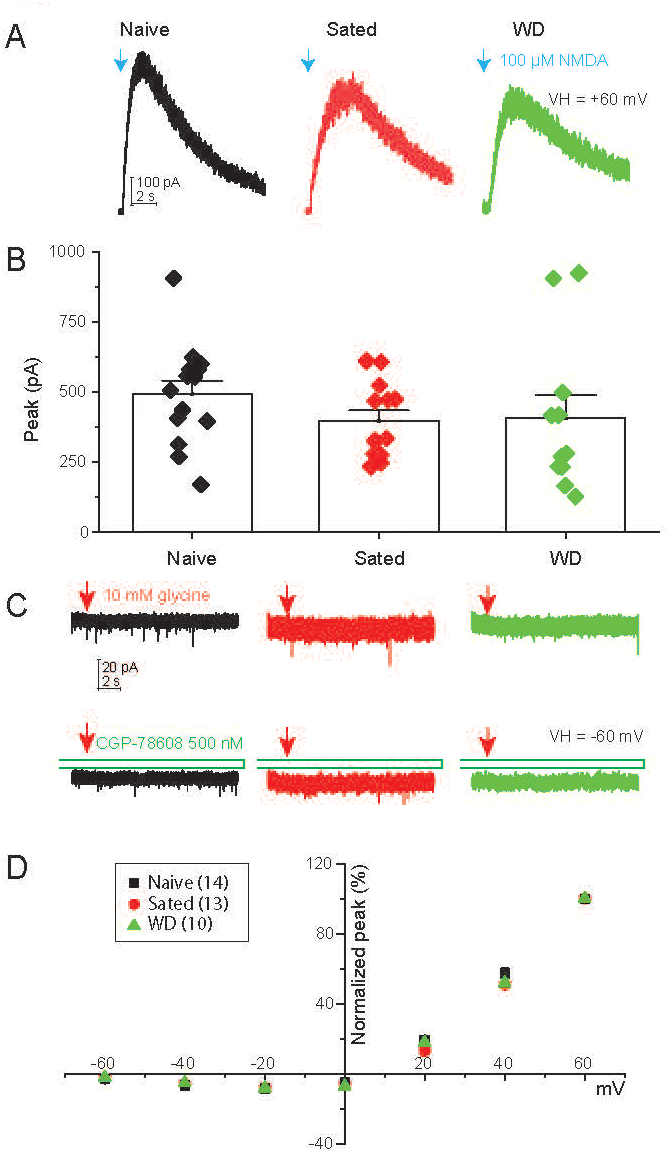
Responses to NMDA and glycine in VTA DA neurons from GluN3A-/- mice. **A.** Typical whole-cell currents induced by puff-applied 100 µM NMDA recorded in VTA DA neurons from naïve, EtOH-sated, and WD mice. Recorded neurons were clamped at +60 mV. Blue arrows indicate NMDA puffs. **B.** The summary of peak NMDA current amplitudes in VTA DA neurons of naïve (493.20 ± 48.52 pA, n=14 cells from 4 males & 5 females), EtOH-sated (398.74 ± 133.73 pA, n=13 cells from 3 males & 4 females), and WD (406.88 ± 82.95 pA, n=11 cells from 3 males & 5 females) mice. No differences were found by one-way ANOVA (F(2,35)= 0.91, p=0.41). **C.** Representative whole-cell currents induced by puffed 10 mM glycine in the absence (top) and in the presence of 500 nM CGP-78608 (bottom) in recorded VTA neurons from naïve (12 cells from 2 males & 3 females), EtOH- sated (13 cells from 3 males & 3 females), and WD (11 cells from 3 males) mice. Recorded neurons were clamped at -60 mV. Opened bars (bottom) indicate the continuous bath application of CGP. The red arrows indicate glycine puffs. **D.** I-V plots of normalized averaged NMDA-induced whole-cell currents in VTA DA neurons of naïve (14 cells from 4 males & 5 females), EtOH-sated (n=13 cells from 3 males & 4 females), and WD (n=10 cells from 3 males & 4 females) mice. All data were normalized to the values obtained at +60 mV (100%). A two-way mixed-design ANOVA found neither differences across EtOH states [F(2,34) = 2.176, p=0.129] nor significant interactions between EtOH states and applied potentials [F(3.108, 52.84=1.687, p=0.179].

In addition to GluN3A, VTA neurons also express NMDARs containing GluN2C/D, which also exhibit reduced Mg²⁺ sensitivity and outward rectification. As previous work has identified cortico-accumbal GluN2C as contributing to EtOH-induced neuroadaptations^75^, these receptors were also examined. Pharmacological experiments with the GluN2C/D potentiator CIQ did not support increased functional contribution of these receptors following chronic EtOH exposure (Supplementary Fig. S1). Together, these findings demonstrate that GluN3A is required for the alcohol-induced remodeling of NMDAR function in VTA dopamine neurons. Furthermore, the absence of glycine-evoked currents in GluN3A-/- mice strongly supports GluN3A, rather than other glycine-responsive NMDAR subtypes such as GluN3B-containing receptors, as the molecular substrate underlying the glycine-sensitive currents observed in WT mice^59^.

## DISCUSSION

The present study identifies functional recruitment of GluN3A-containing NMDARs as a previously unrecognized neuroadaptation produced by chronic alcohol exposure and withdrawal in adult VTA dopamine neurons. Using complementary electrophysiological, pharmacological, and genetic approaches, we demonstrate that chronic EtOH exposure remodels NMDAR signaling by reducing the functional contribution of canonical GluN2A- and GluN2B-containing receptors while promoting the recruitment of functional GluN3A- containing NMDARs. Importantly, the recruited receptors are fully functional, capable of driving membrane depolarization and increasing action potential firing in VTA DA neurons during withdrawal. Thus, rather than representing a quantitative change in NMDAR function alone, our findings reveal a qualitative remodeling of excitatory signaling in which alcohol recruits a receptor subtype with fundamentally different biophysical and signaling properties.

The alcohol-induced impact on NMDAR signaling has been mostly described as changes in canonical GluN2B- and GluN2A-containing NMDARs, both of which contribute to region-specific neuroadaptations and EtOH-related behaviors^46^. For example, EtOH exposure and subsequent withdrawal lead to a persistent increase in the activity of GluN2B-containing NMDARs in the striatum, which promotes further EtOH consumption^44,47^, while increased GluN2A expression has been reported in the cortex and hippocampus^41^ and has been implicated in EtOH tolerance^48^ and dependence^32^. Our findings expand this framework by demonstrating that chronic alcohol exposure also recruits a distinct class of NMDARs that has received little attention in addiction biology.

Although the GluN3A NMDAR subunit was first discovered and characterized in the brain three decades ago^52,53^, its physiological role in the adult brain—and particularly in addiction-related neuroplasticity—remains poorly understood. One reason GluN3A has not received much attention is that its expression is tightly regulated during development, peaking during early postnatal life before declining significantly as neural circuits mature in both humans and in mice^49,52–54^. During development, GluN3A-containing NMDARs regulate synaptic maturation by reducing Ca²⁺ influx and constraining experience-dependent plasticity, thereby limiting excessive NMDAR signaling^49,50,55^.

The present findings suggest that chronic alcohol exposure partially re-engages this developmentally regulated receptor program in the adult VTA. Rather than simply altering the abundance of canonical NMDAR subtypes, our data indicate that alcohol recruits functional GluN3A-containing receptors with reduced Mg²⁺ sensitivity, altered voltage dependence, and lower Ca²⁺ permeability, thereby fundamentally altering the biophysical properties of NMDAR-mediated signaling in VTA DA neurons. Importantly, this interpretation is supported by multiple independent observations, including reduced GluN2A/GluN2B antagonist sensitivity, decreased outward rectification, direct activation by glycine, characteristic GluN3A pharmacology, and complete loss of these adaptations in GluN3A-deficient mice. Interestingly, transient recruitment of GluN3A-containing NMDARs has previously been shown to be required for cocaine-induced synaptic plasticity in VTA DA neurons^61,76^. Together with those findings, our results raise the possibility that recruitment of developmentally regulated GluN3A-containing receptors represents a broader mechanism through which EtOH and addictive drugs remodel mature reward circuits. Whether this reflects a conserved form of drug-induced plasticity or a process unique to specific classes of addictive substances remains an important question for future investigation.

The functional consequences of recruiting GluN3A-containing NMDARs are likely to extend beyond the changes in receptor pharmacology described here. Although functional GluN3A-containing NMDARs were detected in only a small subset of VTA DA neurons from EtOH-naïve mice, their recruitment following chronic EtOH exposure and especially during withdrawal is expected to have important physiological consequences. GluN3A-containing NMDARs differ fundamentally from canonical GluN2-containing receptors because they display reduced Ca²⁺ permeability and diminished voltage-dependent Mg²⁺ block^49,61^. As a consequence, recruitment of these receptors is expected to modify NMDAR-mediated signaling in VTA DA neurons by altering the crucial physiological balance between receptor activation, Ca²⁺ influx, and membrane depolarization.

One physiological process that may be particularly sensitive to these changes is the transition from tonic to burst firing in VTA DA cells, which plays a critical role in reward-based learning, motivation, and memory consolidation^28–30^ and may also contribute to the behavioral manifestations of alcohol withdrawal^77,78^. Activation of canonical NMDARs by glutamate contributes to burst generation because voltage-dependent Mg²⁺ block promotes the membrane potential oscillations that sustain burst firing while limiting depolarization block^79^. Because GluN3A-containing receptors exhibit reduced Mg²⁺ sensitivity, their recruitment during chronic EtOH exposure and withdrawal would be expected to modify these voltage-dependent properties. Although the present study measured spontaneous firing in *ex vivo* VTA slices rather than burst firing directly, the reduced voltage dependence of GluN3A-containing receptors provides a plausible mechanism through which burst generation could be altered during withdrawal. Consistent with this idea, glycine activation of GluN3A-containing NMDARs robustly increased action potential firing in VTA DA neurons, and enhancement of these responses with CGP-78608 frequently drove neurons into depolarization block. These observations therefore support the idea that recruitment of GluN3A-containing receptors can alter the excitability of VTA DA neurons during withdrawal. Indeed, chronic EtOH withdrawal has been reported to reduce spontaneous firing rates, spikes per burst, and absolute burst firing while increasing absolute and relative refractory periods without altering the number of spontaneously active DA neurons^80,81^. Chronic alcohol exposure and withdrawal also produce a persistent hypodopaminergic state characterized by reduced mesolimbic dopamine transmission^82^, a neuroadaptation thought to contribute to withdrawal symptoms and relapse vulnerability. Together with previous evidence that chronic alcohol alters NMDAR-dependent glutamatergic plasticity in reward circuits^5,33^, our findings identify GluN3A recruitment as a plausible cellular mechanism contributing to these physiological adaptations. Whether GluN3A-containing NMDARs directly contribute to withdrawal-associated behaviors or relapse susceptibility remains an important question for future investigation.

The most significant conceptual implication of the present findings is that chronic alcohol exposure may promote neuroadaptations by engaging molecular programs that normally regulate developmental windows of heightened plasticity. During early postnatal development, GluN3A-containing NMDARs contribute to the refinement of neuronal circuits by regulating synaptic maturation and experience-dependent plasticity^49,50,55^.

Developmentally regulated receptor programs are uniquely positioned to modify circuit architecture because they normally govern periods of heightened experience-dependent plasticity during brain maturation. Our observation that chronic EtOH exposure functionally recruits these receptors into adult VTA dopamine neurons raises the possibility that alcohol, rather than simply altering the function of mature glutamate receptors, co-opts GluN3A-related mechanisms to remodel mature reward circuits. This concept is consistent with the current view that addiction is driven by persistent, experience-dependent neuroplasticity, whereby repeated drug exposure produces enduring modifications of neural circuits that stabilize drug-associated memories and increase vulnerability to relapse^6,8,83,84^.

In addition to altering NMDAR signaling, recruitment of GluN3A-containing receptors fundamentally changes the physiological role of glycine within the adult VTA. Whereas glycine normally functions as a co-agonist at canonical NMDARs, incorporation of GluN3A allows glycine to directly activate excitatory NMDAR currents^56,57,59–61^—even while activities of conventional NMDA-gated NMDARs are partially blocked by Mg^2+^. Our findings demonstrate that withdrawal from chronic EtOH exposure markedly increases the prevalence of these glycine-responsive receptors and that glycine alone is sufficient to depolarize VTA DA neurons and increase action potential firing. Whether endogenous glycine concentrations during alcohol withdrawal are sufficient to engage these receptors under physiological conditions remains an important question for future investigation.

In conclusion, our findings broaden the current view of alcohol-induced glutamatergic plasticity by demonstrating that chronic EtOH exposure remodels NMDAR signaling through recruitment of a developmentally regulated receptor subtype rather than solely through modifications of canonical GluN2-containing receptors. This observation suggests that alcohol-induced neuroadaptations involve qualitative changes in receptor identity that fundamentally alter the properties of NMDAR signaling in adult VTA dopamine neurons. Notably, because GluN3A-containing receptors remain largely unexplored in AUD, they represent a potentially novel therapeutic target for limiting maladaptive glutamatergic plasticity associated with chronic alcohol exposure and withdrawal. Together, these findings raise the possibility that recruitment of a NMDAR subtype predominantly expressed during development represents a previously unrecognized mechanism contributing to the persistent circuit remodeling that characterizes EtOH addiction.

## ACKNOWLEDGMENTS

This work was supported by the National Institute on Alcohol Abuse and Alcoholism, grant UO1 AA025931 and a Bench to Bedside (BtB) supplement to UO1 AA025931 (MDB) and the National Institute on Drug Abuse, R37 DA053296 (JAD). The Dani Lab is also supported by a generous award from the Chernowitz Medical Research Foundation.

