## Supplemental data for "Chronic Alcohol Exposure Alters Expression of GluN3A-NMDA Receptors in Adult VTA DA Neurons"

### Chronic EtOH exposure does not affect GluN2C/2D-containing NMDA receptors.

Because tri-heteromeric NMDARs containing the GluN2C/D subunit are expressed on VTA neurons, exhibit reduced outward rectification and could be involved in the modulation of reward signals[1], we examined whether chronic EtOH exposure and withdrawal could also increase the expression of GluN2C/D-containing NMDARs. Considering that the effects of EtOH are most prominent during withdrawal, we compared NMDA-gated currents in slices from EtOH-naïve and EtOH-withdrawn mice in the presence of CIQ, which was bath-applied at a 20 μM concentration. CIQ is a positive allosteric modulator that selectively potentiates NMDARs containing GluN2C/2D but has no effect on NMDARs that contain exclusively GluN1/GluN2A/2B subunits [2,3].

Mice were exposed to the drinking-in-the-dark paradigm—a widely-used model of binge-like ethanol drinking [4-6] with acclimation to the 50ml sipper tube in the home-cage lasting for at least 1 week. During the experimental phase, the sipper tube was replaced with either 15% (v/v) unsweetened ethanol or filtered tap water (control), beginning approximately 2-h into the dark phase of the light cycle. Mice were allowed to drink for 4-h, whereupon the sipper tube was replaced with filtered tap water. Sessions occurred daily and continued for 5-8 weeks. Brains were harvested for slice electrophysiology ~20-h after ethanol withdrawal.

CIQ potentiated NMDAR currents in both EtOH naïve and EtOH withdrawn mice (Fig. S1 A-D), but there was no significant difference between the two experimental groups (Fig. S1E, t-test, $p=0.38$) indicating that, as previously reported [1], tri-heteromeric GluN2C/2D NMDARs are expressed by VTA neurons in physiological, EtOH-naïve conditions and that expression of those receptors is not altered by withdrawal from chronic EtOH exposure.

Supplemental Figure 1

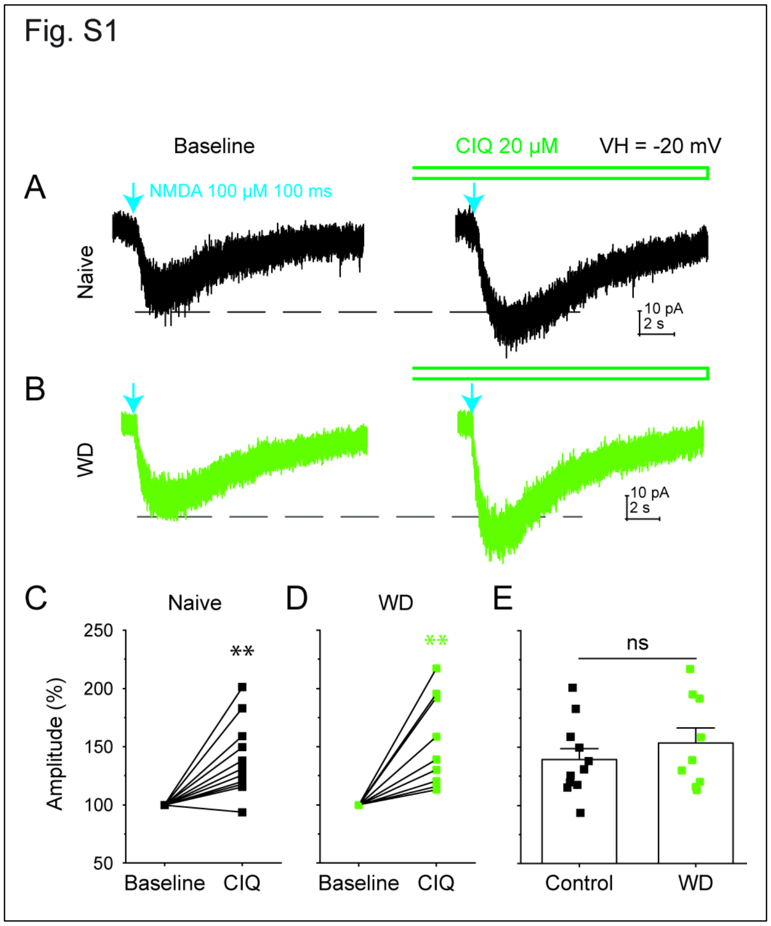


**Figure S1. Potentiation effects of 20 µM CIQ on NMDA currents in VTA DA neurons**

**A-B**. Representative NMDA currents induced by 100 µM NMDA puffs before (Baseline, Left) and after bath application of CIQ (Right) in VTA DA neurons of control (**A**) and EtOH WD (**B**) mice. The blue arrows indicate NMDA puffs. The open bars indicate bath application of 20 µM CIQ. **C-D.** Comparison of normalized peak amplitude of NMDA currents before and after CIQ in control (C, n=11 cells from 4 males & 3 females) and EtOH WD (D, n=9 cells from 3 males & 3 females) mice. CIQ significantly increased amplitude in both control [$t(10)=4.14$, $p<0.005$] and WD mice [$t(8)=4.1$, $p<0.005$] relative to baseline (100%). **E**. Bar graphs summarize and compare the potentiation of NMDA currents by CIQ in control and EtOH WD mice. There was no significant difference in the magnitude of the response to CIQ between control and WD mice, $t(18)=-0.896$, $p=0.382$. **$p<0.01$.
